# The metronomic combination of paclitaxel with cholinergic agonists inhibits triple negative breast tumor progression. Participation of M_2_ receptor subtype

**DOI:** 10.1101/858969

**Authors:** Alejandro J. Español, Agustina Salem, María Di Bari, Ilaria Cristofaro, Yamila Sanchez, Ada M. Tata, María E. Sales

**Affiliations:** Center of Pharmacological and Botanical Studies (CEFYBO). CONICET. Buenos Aires, Argentine; Second Department of Pharmacology, School of Medicine, University of Buenos Aires. Buenos Aires, Argentine; Department of Biology and Biotechnologies Charles Darwin, Sapienza University of Rome. Rome, Italy; Center of Neurobiology Daniel Bovet, Sapienza University of Rome. Rome, Italy

## Abstract

Triple negative tumors are more aggressive than other breast cancer subtypes and there is a lack of specific therapeutic targets on them. Since muscarinic receptors have been linked to tumor progression, we investigated the effect of metronomic therapy employing a traditional anti-cancer drug, paclitaxel plus muscarinic agonists at low doses on this type of tumor. We observed that MDA-MB231 tumor cells express muscarinic receptors, while they are absent in the non-tumorigenic MCF-10A cell line, which was used as control. The addition of carbachol or arecaidine propargyl ester, a non-selective or a selective subtype 2 muscarinic (M_2_) receptor agonist respectively, plus paclitaxel reduces cell viability involving a down-regulation in the expression of ATP “binding cassette” G2 drug transporter and epidermal growth factor receptor. We also detected an inhibition of tumor cell migration and anti-angiogenic effects produced by those drug combinations *in vitro* and *in vivo* (in NUDE mice) respectively. Our findings provide substantial evidence about M_2_ receptors as therapeutic target for the treatment of triple negative tumors.

## Introduction

Breast cancer is yet the most frequent type of malignancy in women and represents a major and unsolved problem for public health [1]. Luminal and triple negative (TN) represent the two opposite ends of the molecular classification of breast tumors and they thoroughly differ regarding treatment and patients’survival [2]. The TN tumors are typically larger in size, higher grade than other breast cancers, and they also exhibit an aggressive clinical behavior, frequently resulting in early metastatic dissemination, particularly to visceral sites. As a result of these characteristics, TN breast cancers are associated with poor prognosis in comparison to luminal breast tumors [3]. Considering the treatment of TN tumors, classical modalities have improved the overall outlook and quality of life for women with this type of breast cancer. However, because of recurrence and/or the development of resistance to cytotoxic drugs administered to patients, produced by a complex mechanism mediated by different types of proteins such as ATP “binding cassette” (ABC) transporters, a considerable amount of patients still succumb to this disease highlighting the need to find new therapeutic approaches [4]. Regarding the latter, the usage of low dose chemotherapy with short drug free intervals, named metronomic therapy has emerged as a novel regimen for cancer treatment [5]. It exerts very low incidence of side effects and could add new beneficial actions on immune system and tumor microenvironment [6]. This new strategy also needs the identification of new therapeutic targets to improve the benefits for breast cancer patients.

Non-neuronal cholinergic system (nNCS) has been involved either in physiological or in pathological processes. The nNCS is formed by acethylcholyne (ACh), the enzymes that synthesize and degrade ACh and cholinergic receptors expressed in non-neuronal cells. Muscarinic receptors belong to this group of proteins and have been involved in the progression of different type of tumors such as lung, colon and prostate [7-9]. We demonstrated that muscarinic receptors are expressed in tumor samples from patients with breast cancer in different stages and also in human MCF-7 cells derived from a luminal, estrogen-dependent adenocarcinoma, the most frequent type of breast tumor in women [10,11]. Muscarinic receptors belong to the G-protein coupled receptors family which constitutes the largest family of cell surface receptors involved in signal transduction. Five subtypes have been identified by molecular cloning: M_1_-M_5_. Their role in the regulation of important cell functions like mitosis, cell morphology, locomotion and immune response which are key steps during tumor progression has been documented [12]. The long-term activation of these receptors with the agonist carbachol stimulates cytotoxicity either in human or in murine breast tumor cells [13,14]. In the last years, several reports demonstrated that the activation of M_2_ receptor subtype by a selective agonist was able to arrest cell proliferation in different tumor cell lines [15-17]. Moreover, M_2_ receptor activation reduced cell survival, inducing oxidative stress and severe apoptosis in malignant cells derived from human glioblastoma [17]. The MDA-MB231 is a human cell line derived from a TN breast tumor, which does not express estrogen/progesterone receptors or HER2 protein.

The aim of our work is to investigate the ability of a combination of low doses of paclitaxel (PX) and a muscarinic agonist to inhibit different steps of TN breast tumor progression. In this work, we identified different subtypes of muscarinic receptors in MDA-MB231 cells by Western blot, and demonstrated that the combination of PX plus carbachol or arecaidine propargyl ester (APE), a non selective or a M_2_ selective agonist, reduced cell viability, migration, vascular endothelial growth factor-A (VEGF-A) expression as well as *in vivo* angiogenesis. We also observed a down-regulation in the expression of ABCG2 transporter and epidermal growth factor receptor (EGFR) in tumor cells by Western blot, after PX plus carbachol or APE administration revealing that both proteins could be involved in the mechanism of action of this treatment.

## Materials and Methods

### Cell culture

The human breast adenocarcinoma cell line MDA-MB231 (CRM-HTB-26) and MDA-MB468 (HTB-132) were obtained from the American Type Culture Collection (ATCC; Manassas, USA) and cultured in DMEM (Invitrogen Inc., Carlsbad, USA) with 2 mM L-glutamine and 80 μg/ml gentamycin, supplemented with 10% heat inactivated FBS (Internegocios SA, Mercedes, Argentine) at 37°C in a humidified 5% CO_2_ air. The MCF-10A cells (CRL-10317) were also purchased by ATCC and constitute a non-tumorigenic cell line derived from human mammary tissue. These cells were grown on tissue culture plastic dishes in DMEM:F12(1:1) (Invitrogen Inc., Carlsbad, USA) supplemented with 10% FBS, hydrocortisone (0.5 μg/ml), insulin (10 μg/ml), and hEGF (20 ng/ml). Cell lines were detached using the following buffer: 0.25% trypsin and 0.02% EDTA in Ca^2+^ and Mg^2+^ free PBS from confluent monolayers. The medium was replaced three times a week. Cell viability was assayed by Trypan blue exclusion test and the absence of mycoplasma was confirmed by Hoechst staining [18].

### Detection of muscarinic receptors by Western blot

Cells (2×10^6^) were washed twice with PBS and lysed in 1 ml of 50 mM Tris-HCl, 50 mM NaCl, 5 mM NaF, 5 mM MgCl_2_, 1 mM EDTA, 1 mM EGTA, 5 mM phenylmethanesulfonyl fluoride, 1% Triton X-100 and 10 μg/ml trypsin inhibitor, aprotinin and leupeptin, pH 7.4. After 1 h in an ice bath, lysates were centrifuged at 800 g for 20 min at 4°C. The supernatants were stored at −80°C and protein concentration was determined by the method of Bradford [19]. Samples (80 μg protein per lane) were subjected to 10% SDS-PAGE minigel electrophoresis, transferred to nitrocellulose membranes, and incubated overnight with goat anti-human M_1_, M_2_ or M_3_ receptor polyclonal antibodies or rabbit anti-human M_4_ or M_5_ receptor polyclonal antibodies (Santa Cruz Biotechnology Inc., USA) all diluted 1:200. Then strips were incubated with anti-rabbit IgG or anti-goat IgG coupled to horseradish peroxidase, both diluted 1:10000 in 20 mM Tris-HCl buffer, 150 mM NaCl and 0.05% Tween 20 (TBS-T) at 37°C for 1 h. Bands were visualized by chemiluminescence. The results of the densitometric analysis of the bands were expressed as optical density (O.D.) units relative to the expression of glyceraldehyde 3-phosphate dehydrogenase (GAPDH) (Santa Cruz Biotechnology Inc., USA) [20].

### Cell viability assay

The inhibition in cell viability exerted by different treatments for one or three cycles of 48h was evaluated by using the soluble tetrazolium salt 3-(4,5-dimethylthiazol-2-yl)-2,5-diphenyltetrazolium bromide (MTT) colorimetric assay (Life Technologies, Eugene, USA). In living cells, MTT is reduced to formazan. Cells were seeded in 96-well plates at a density of 4×10^3^cells per well in culture medium supplemented with 5% FBS and then left to adhere overnight. When cells reached 60-70% of confluence they were deprived of FBS 24 h previous to the assay to induce the synchronization of cultures. Then, cells were treated with PX (Bristol-Myers Squibb, Vicente López, Argentine), carbachol or arecaidine propargyl ester (APE) (non-selective or selective M_2_ receptor agonist respectively) alone or in combination in medium supplemented with 2% FBS, during 48 h in triplicate. Also the effect of doxorubicin (Glenmark Generics S.A., Pilar, Argentine) alone or combined with muscarinic agonists was analyzed. To inhibit the action of cholinergic agonists, cells were previously treated with atropine at 10^−9^M or methoctramine at 10^−5^M (non-selective or M_2_ receptor selective antagonist respectively).

After treatment, to detect viable cells, the medium was replaced by 110 μl of MTT solution that was prepared by diluting 10 μl of 5 mg/ml MTT in PBS, in 100 μl medium free of phenol red and FBS to each well. After incubation for 4 h at 37°C, the production of formazan was evaluated by measuring the absorbance at 540 nm with an ELISA reader (BioTek, Winooski, USA). Values are mean ± S.E.M and results are expressed as the percentage of inhibition in cell viability in comparison to control (cells without treatment).

### ATP binding cassette transporter G2, epidermal growth factor receptor and vascular endothelial growth factor-A detection by Western blot

Cells (2×10^6^) were treated for three cycles of 48h with the treatments and samples were prepared as it was indicated to detect muscarinic receptors. Then, samples (80 μg protein per lane) were subjected to 8-10% SDS-PAGE minigel electrophoresis, transferred to nitrocellulose membranes, and incubated overnight with a rabbit anti-human ABCG2 polyclonal antibody (Santa Cruz Biotechnology Inc., Dallas, USA) diluted 1:200, a rabbit anti-human EGFR monoclonal antibody (EMD Millipore-MERCK, Temecula, USA) diluted 1:1000 or a rabbit anti-human VEGF-A polyclonal antibody (Abcam, Eugene, USA) diluted 1:1000. Then strips were incubated with horseradish peroxidase-linked anti-rabbit IgG, diluted 1:10000 in TBS-T at 37°C for 1 h. Bands were visualized by chemiluminiscence. Quantification of the bands was performed by densitometric analysis using Image J program (NIH) and was expressed as O.D. units in comparison to the expression of GAPDH that was used as loading control [21].

### Cell migration assay

To quantify the ability of tumor cells to migrate, an *in vitro* wound healing assay was performed according to previously described methods [22]. Cells (1.5×10^5^/well) were seeded in 24-well plates with 0.5 ml of DMEM medium and left to adhere. Then, the cell monolayer was scratched with 200 µl pipette tip, washed twice with PBS and fresh culture medium containing different drugs was added. The migration of cells was photographed at regular intervals from the beginning of the assay during 28 h and the uncovered area was integrated with image J software (NIH). The results were expressed as the percentage of covered area.

### Tumor-induced angiogenesis

Female NUDE mice (3 months old and 25 g/mouse) were purchased from the animal facility of the Faculty of Veterinary Sciences, La Plata National University (La Plata, Argentine). Animals were kept under specific pathogen-free conditions following the protocol designed by the National Institute of Health (NIH, USA) in the Guidelines for the care and handling of laboratory animals (1986). Experimental procedures were approved by the Institutional Committee for the Care and Use of Laboratory Animals (CICUAL) from the School of Medicine, University of Buenos Aires (Protocol number 20020170100227BA).

Neovascularization induced by tumor cells, was analyzed using an *in vivo* bioassay previously described [23]. Briefly, cell suspension was prepared by detaching and washing MDA-MB231 cells in DMEM, the concentration was adjusted to 3×10^6^ cells/ml and 0.1 ml containing 3×10^5^ cells were injected intradermically in each flank of female NUDE mice. Animals were treated intraperitoneally (i.p.) 24 h post-injection of cells, with carbachol (0.94 pg/mouse) or APE (4.66 µg/mouse) alone or in combination with PX (0.51 ng/mouse). Treatments were administered in 3 doses with 48 h intervals. Atropine (0.17 ng/mouse) or methoctramine (3.5 µg/mouse) were inoculated i.p. 20 min before other drugs. Drugs were dissolved in PBS under sterile conditions. Doses were calculated considering the concentrations added to cells *in vitro*, the amount of inoculated cells and the lasting of treatment. Each experiment includes 10 different treatments with 3 animals per treatment/group repeated 3 times.

On day 6, animals were sacrificed by CO_2_ inhalation and the skin was exposed. The vascular response was observed in the inner surface of the skin with a dissecting microscope (Konus Corporation, Miami, USA) at a 6.4 X magnification, and the sites of inoculation were photographed with an incorporated digital camera (Canon Power Shot A75, Canon Inc., Lake Success, USA). Photographs were projected onto a reticular screen to count the number of vessels per mm^2^ of skin. Angiogenesis was quantified as vessel density, determined by the formula: Σ number of vessels in each square/total number of squares [23].

### EC50 calculation

Using the GraphPad Prism 6 program, dose–response data were transformed, changed to percentage and fitted using to a sigmoidal curve following a maximal effective concentration (Emax) model with at least six data points. The EC50 and Emax values were obtained from this analysis. Only data with less than 20% in the coefficient of variation for EC50 values were considered.

### Statistical Analysis

Results were expressed as mean ± S.E.M. The GraphPad Prism6 computer program was used employing one-way ANOVA analysis for paired samples to obtain the significance of differences between mean values in all control and test samples. The analysis was complemented by using a Tukey test to compare among mean values. Differences between means were considered significant if P<0.05. The data and statistical analysis complied with the recommendations on experimental design and analysis in pharmacology [24]. When it was necessary, values were also analyzed using Chou Talalay method to evaluate drug combination effects [25].

### Drugs

All drugs were purchased from Sigma Chemical Co. (St. Louis, USA) unless otherwise stated. Solutions were prepared fresh daily.

## Results

### Muscarinic agonists modify MDA-MB231 cell viability

In this work, we demonstrated by Western blot the expression of M_1_, M_2_, M_4_ and M_5_ receptors in MDA-MB231 cell lysates (Fig 1A). The non-tumorigenic mammary cell line MCF-10A lacks all subtypes of muscarinic receptors (Fig 1B).

**Fig 1.**
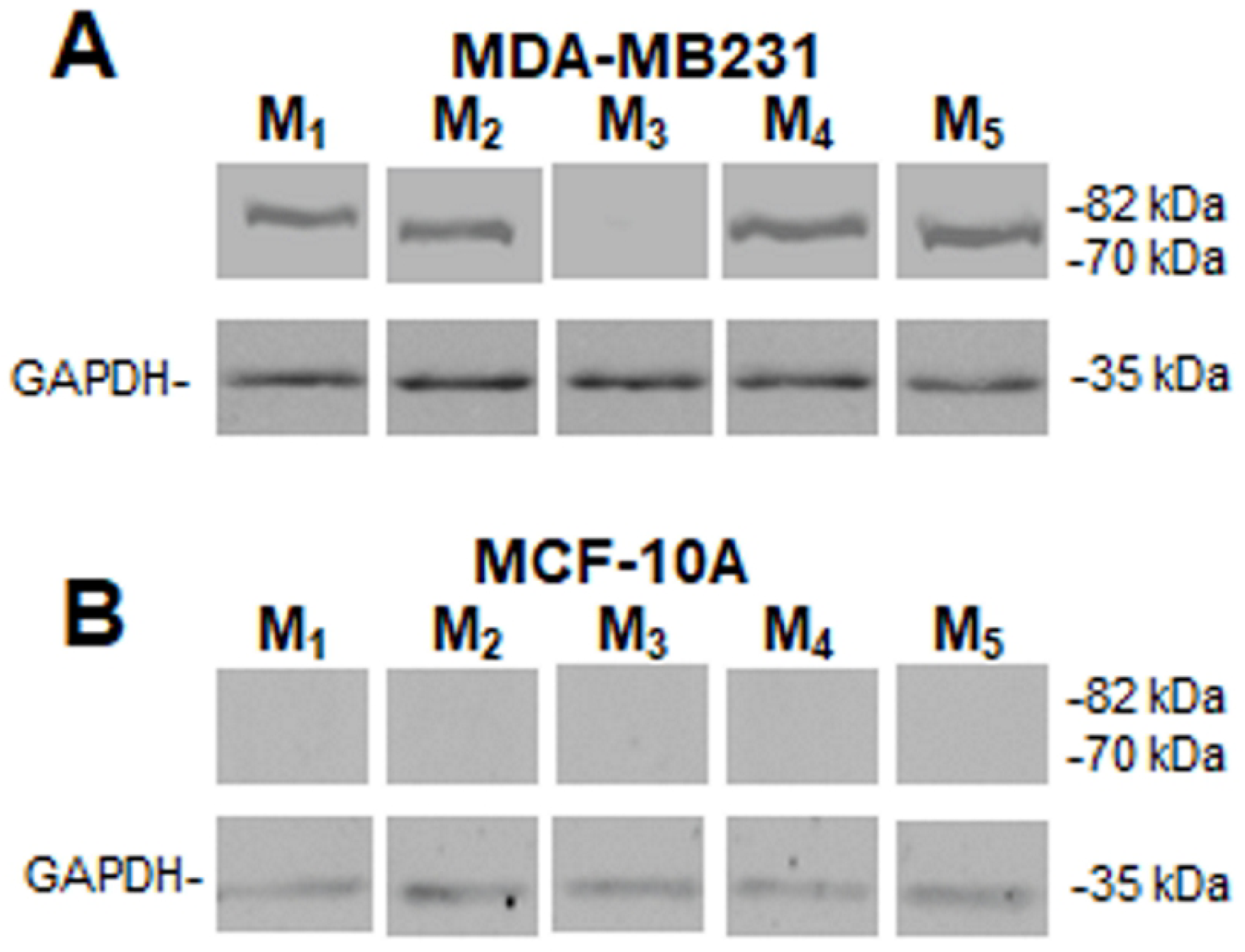
Muscarinic receptors’ expression. Western blot assay to detect muscarinic (M) receptor subtypes in A) MDA-MB231cells or B) in MCF-10A cells. Molecular weights are indicated on the right. The expression of glyceraldehyde 3-phosphate dehydrogenase (GAPDH) protein was used as loading control. One representative experiment of 3 is shown.

In addition, we analyzed the effect of increasing concentrations of the non-selective muscarinic agonist carbachol added to MDA-MB231 cells in culture. Figure 2A shows that carbachol produced an inhibition in cell viability in a concentration-dependent manner (EC50: 1.2×10^−11^M). Also APE was effective to reduce tumor cell viability at concentrations higher than 10^−5^M (EC50: 3.1×10^−5^M) (Fig 2B). Because of the absence of muscarinic receptors in non-tumorigenic MCF-10A cells, neither carbachol nor APE modified cell viability at any concentration tested (Figs 2A and 2B). We confirmed the cytotoxic activity of PX on breast cancer cells. In our system, this chemotherapeutic agent decreased viability at concentrations ≥ 10^−8^M in tumor cells (EC50: 8.5×10^−7^M). As an undesirable action, PX also reduced MCF-10A cell viability from10^−7^M (Fig 2C).

**Fig 2.**
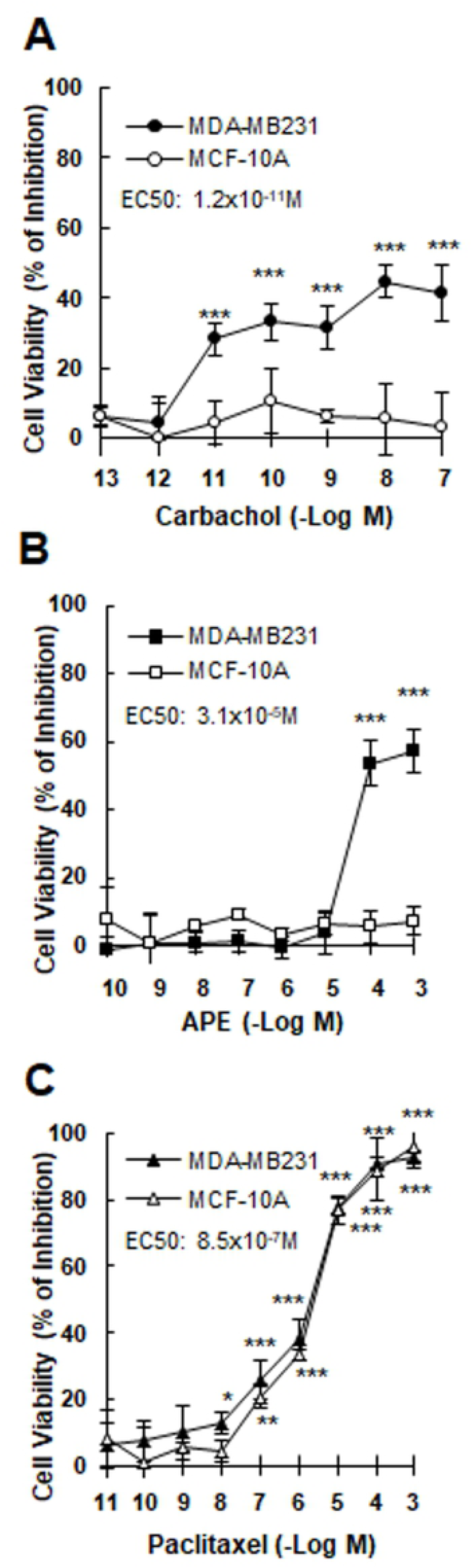
Effect of muscarinic agonists or paclitaxel on breast cell viability. Concentration-response curves of A) carbachol, B) arecaidine propargyl ester (APE) or C) paclitaxel on MDA-MB231 cells or MCF-10A cells. Results were expressed as percent of inhibition in cell viability respect to control (cells without treatment). EC50: effective concentration that produces half maximal response. Values are mean ± S.E.M. of 5 experiments performed in duplicate. (*P<0.05; **P<0.001;***P<0.0001 vs. control: untreated cells).

### Effects produced by the combined treatment of paclitaxel with muscarinic agonists on MDA-MB231 cells

In order to determine the ability of muscarinic agonists to synergize the action of PX on tumor cells, we performed concentration-response curves of this drug in the presence of the EC25 of carbachol or APE (8.6×10^−12^M or 1.1×10^−5^M respectively) to evaluate cell viability (Fig 3). The addition of carbachol shifted to the left the dose-response curve of PX modifying the EC50 value by more than one order of magnitude (EC50 PX: 8.5×10^−7^M; EC50PX+carbachol: 1.1×10^−8^M) (Fig. 3A). Similar results were obtained when APE was added to the concentration-response curve of PX (EC50: PX+APE: 3.5×10^−8^M) on tumor cells (Fig 3B). These results and the combination index (CI)<1 calculated by the method of Chou Talalay using CompuSyn Software (PX+carbachol: 0.10077; PX+APE: 0.13951) indicate a synergism of potentiation for both pairs of drugs.

**Fig 3.**
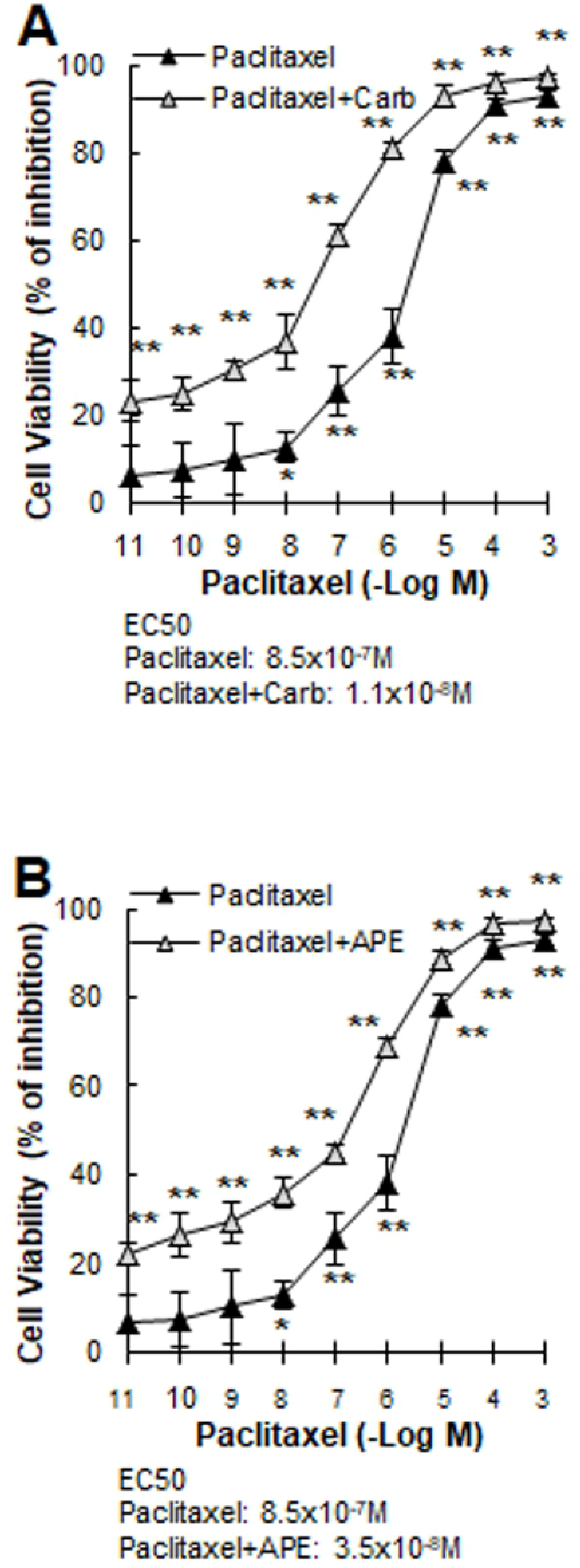
Effect of muscarinic agonists on the concentration-response curve of paclitaxel. The MDA-MB231 cells were treated with increasing concentrations of paclitaxel in the absence or presence of A) carbachol (8.6×10^−12^M) or B) arecaidine propargyl ester (APE) (1.1×10^−5^M) during 48 h. Results were expressed as percent of inhibition in cell viability respect to control (cells without treatment). EC50: effective concentration that produces half maximal response. Values are mean ± S.E.M. of 5 experiments performed in duplicate. (*P<0.05; **P<0.0001 vs. control: untreated cells).

Taking into account the undesirable effect induced by chemotherapy at usual doses (≥10^−6^M), we analyzed the ability of PX at the first effective concentration (10^−8^M) combined with the EC25 of carbachol or APE on the viability of tumor cells in order to mimic the dosage of metronomic therapy (Fig 4). The combination of PX plus carbachol significantly reduced cell viability of MDA-MB231 cells. The latter effect was prevented in the presence of 10^−9^M atropine, a non-selective muscarinic antagonist. This combination did not exert any action on non-tumorigenic MCF-10A mammary cells (Fig 4A). Taking into account that M_2_ receptors are expressed in tumor cells, we analyzed the effect of the M_2_ selective agonist APE in combination with PX. The addition of APE (1.1×10^−5^M) combined with PX potentiated the effect of PX alone to reduce tumor cell viability (Fig 4B). The action of APE plus PX was reverted by the previous addition of the M_2_ selective antagonist methoctramine (10^−5^M) revealing the main participation of this receptor subtype in the reduction of tumor cell viability (Fig 4C). We also proved that the addition of carbachol, APE and/or PX at the same concentrations did not modify MCF-10A cell viability (Figs 4A and 4B).

**Fig 4.**
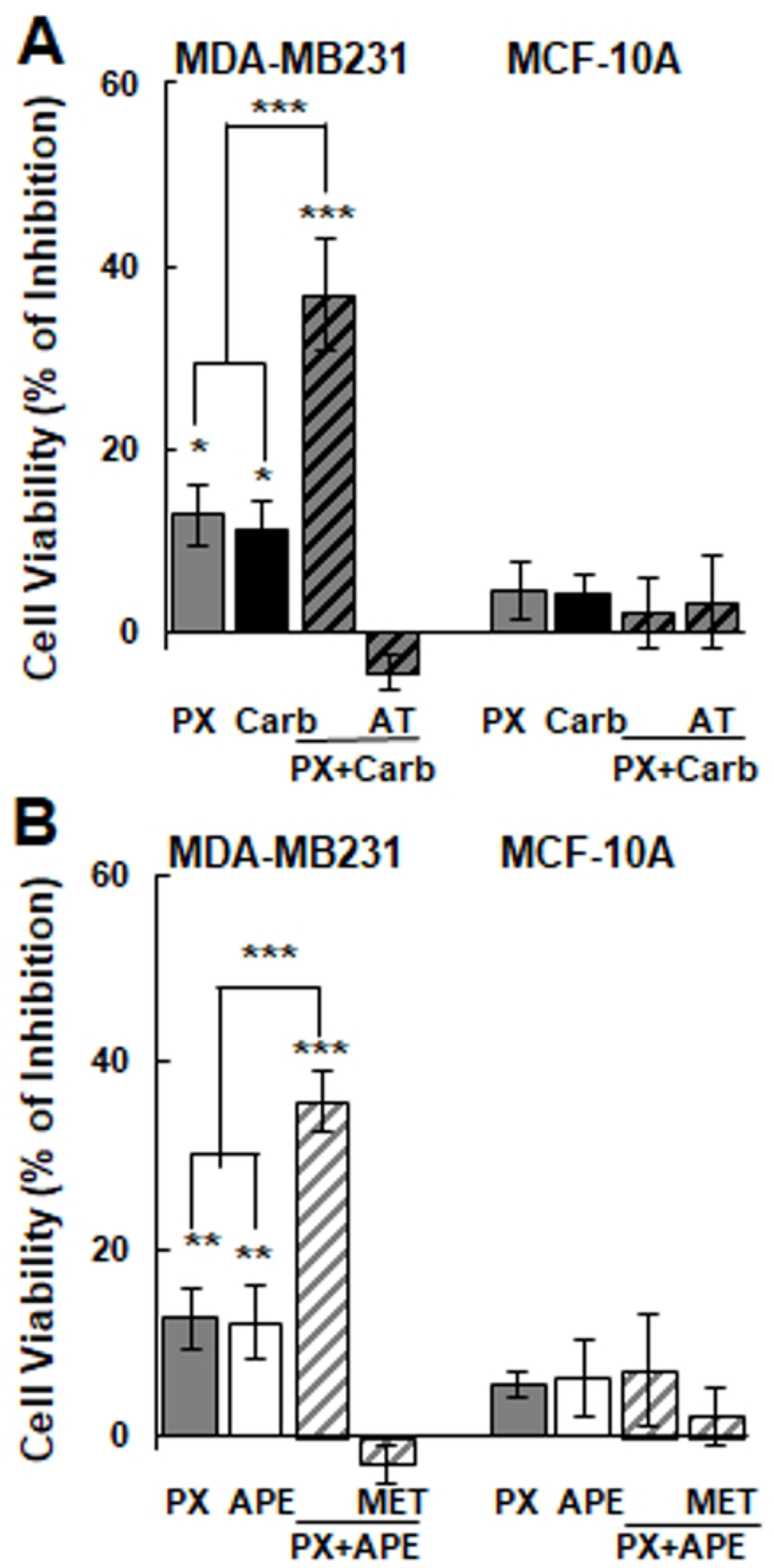
Effect of the combination of paclitaxel with a muscarinic agonist on breast cell viability. Cells were treated with A) paclitaxel (PX) (10^−8^M) combined with carbachol (Carb) (8.6×10^−12^M) in the absence or presence of atropine (AT) (10^−9^M) or B) PX (10^−8^M) was combined with arecaidine propargyl ester (APE) (1.1×10^−5^M) in the absence or presence of methoctramine (MET) (10^−5^M). Results are expressed as percent of inhibition in cell viability respect to control (cells without treatment). Values are mean ± S.E.M. of 5 experiments performed in duplicate. (*P<0.01; **P<0.001; ***P<0.0001 vs. control, PX or Carb).

To confirm that muscarinic agonists can synergize the action of another cytotoxic drug reducing tumor cell viability, we tested the effect of 10^−8^M doxorubicin (the first effective concentration) frequently used in breast cancer treatment combined with carbachol or APE both at EC25 on MDA-MB231 cells (Table 1).

**Table 1.** Effect of the combination of doxorubicin with a muscarinic agonist on MDA-MB231 cell viability. Cells were treated with doxorubicin (DX) (10^−8^M) combined with carbachol (Carb) (8.6×10^−12^ M) or with arecaidine propargyl ester (APE) (1.1×10^−5^ M) during 48 h in the absence or presence of atropine (AT) (10^−9^M) or methoctramine (MET) (10^−5^M) respectively. Results are expressed as cell viability (% of inhibition) respect to control (cells without treatment). Values are mean ± S.E.M. of 5 experiments performed in duplicate. *P<0.001; **P<0.0001 vs. control (cells without treatment).

| MDA-MB231 |  |
| --- | --- |
| Treatment | Cell Viability (% of Inhibition) |
| DX + Carb | **35.3±0.8 \*\*** |
| DX + Carb + AT | -6.7±2.1 |
| Carb | 11.2±3.2 * |
| DX | 12.4±4.6 * |
| DX + APE | **33.3±2.1 \*\*** |
| DX + APE + MET | -7.7±2.6 |
| APE | 12.2±3.9 * |

Either the presence of carbachol or APE potentiated the effect of doxorubicin by reducing tumor cell viability. The effect was prevented by preincubating cells with atropine or methoctramine. Moreover, in MDA-MB468 cells derived from another subtype of TN tumor, we observed that the addition of carbachol or APE at the EC25 (1.1×10^−10^M or 1.3×10^−7^M respectively) to the first effective concentration of PX (10^−9^M) was also effective to reduce cell viability. These tumor cells also express all subtypes of muscarinic receptors (S1 Fig). The addition of atropine or methoctramine before the combined treatment significantly reduced this effect (Table 2).

**Table 2.**
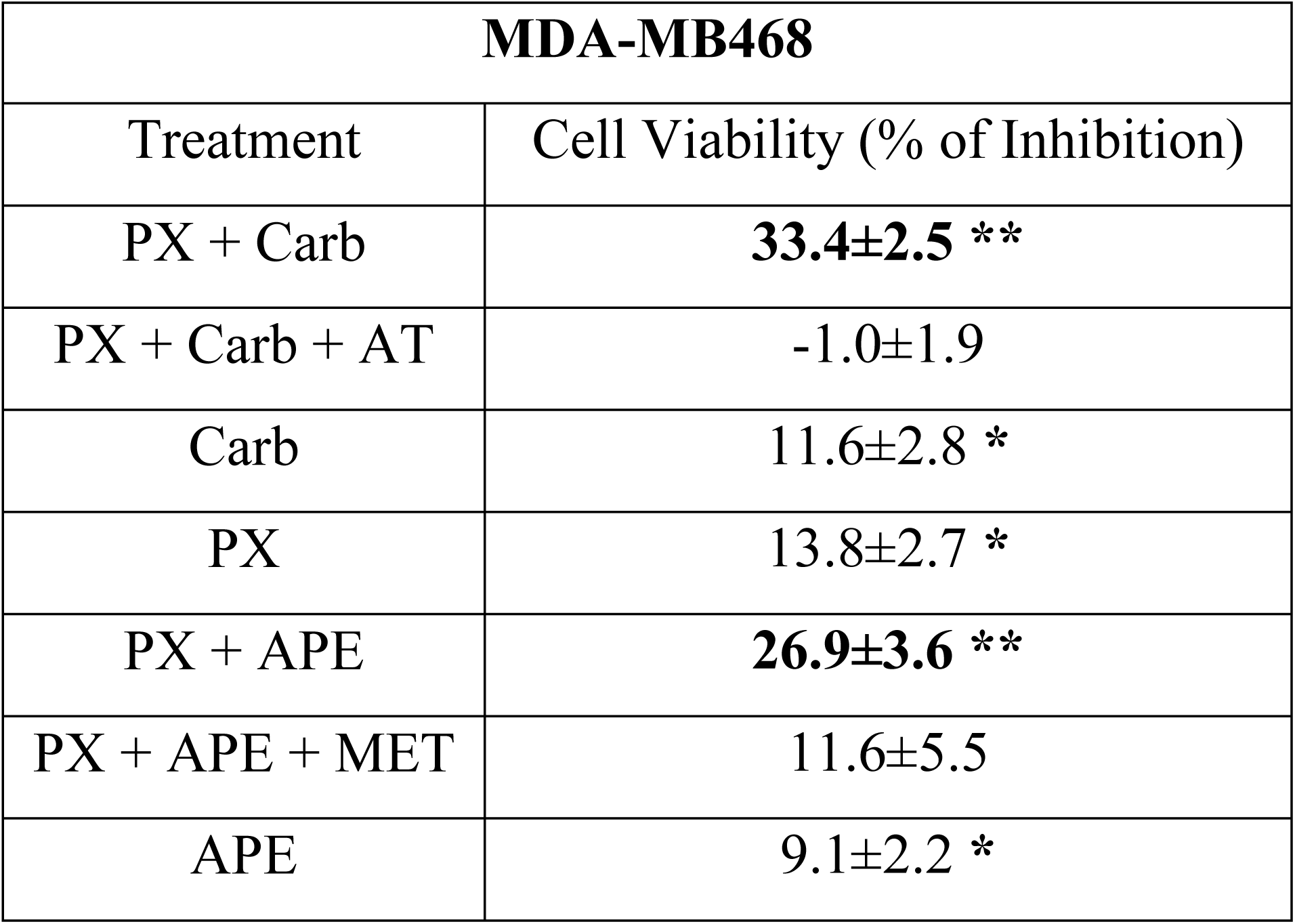
Effect of the combination of paclitaxel with a muscarinic agonist on MDA-MB468 cell viability. Cells were treated with paclitaxel (PX) (10^−9^M) combined with carbachol (Carb) (1.2×10^−10^ M) or with arecaidine propargyl ester (APE) (1.4×10^−7^M) during 48 h in the absence or presence of atropine (AT) (10^−9^M) or methoctramine (MET) (10^−5^M) respectively. Results are expressed as cell viability (% of inhibition) respect to control (cells without treatment). Values are mean ± S.E.M. of 5 experiments performed in duplicate. *P<0.001; **P<0.0001 vs. control (cells without treatment).

| MDA-MB468 |  |
| --- | --- |
| Treatment | Cell Viability (% of Inhibition) |
| PX + Carb | **33.4±2.5 \*\*** |
| PX + Carb + AT | -1.0±1.9 |
| Carb | 11.6±2.8 * |
| PX | 13.8±2.7 * |
| PX + APE | **26.9±3.6 \*\*** |
| PX + APE + MET | 11.6±5.5 |
| APE | 9.1±2.2 * |

We confirm that the combination of PX at the first effective concentration plus carbachol or APE at the EC25 for three cycles potently reduced cell viability of MDA-MB231 cells by more than 60%. This effect was prevented by the previous addition of atropine o methoctramine respectively (Figs 5A and 5B).

**Fig 5.**
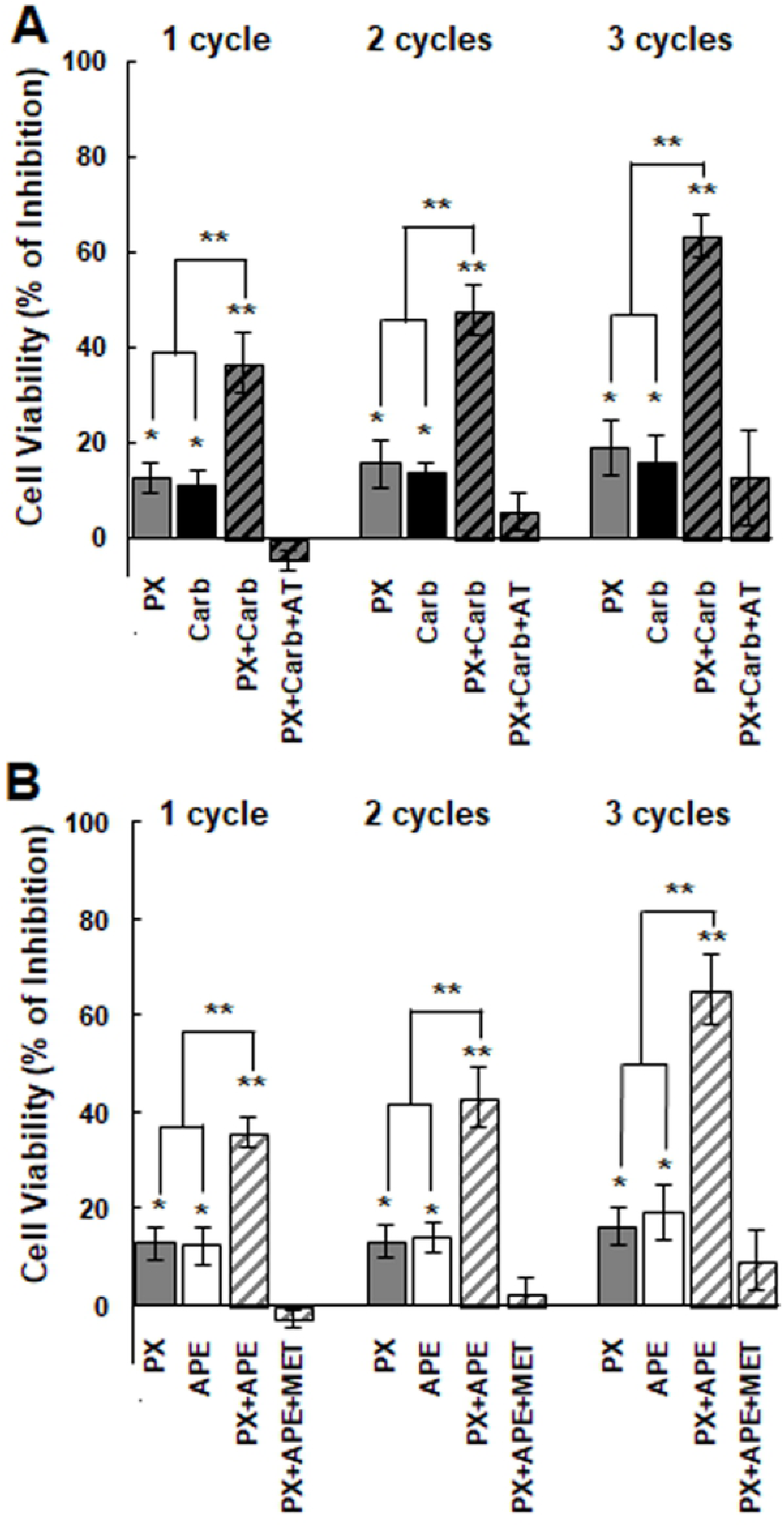
Effect of the combination of paclitaxel with a muscarinic agonist on breast cell viability administered in three cycles. Cells were treated with A) paclitaxel (PX) (10^−8^M) combined with carbachol (Carb) (8.6×10^−12^M) in the absence or presence of atropine (AT) (10^−9^M) or B) PX (10^−8^M) was combined with arecaidine propargyl ester (APE) (1.1×10^−5^M) in the absence or presence of methoctramine (MET) (10^−5^M). Results are expressed as percent of inhibition in cell viability respect to control (cells without treatment). Values are mean ± S.E.M. of 4 experiments performed in duplicate. (*P<0.05; **P<0.01***P<0.001; ****P<0.0001 vs. control, PX or Carb).

In order to investigate the mechanism of action involved in the effect produced by the combinations, we analyzed the expression of ABCG2 and EGFR in tumor cells. The addition of PX with carbachol (Fig 6A) or APE (Fig 6B) significantly reduced the expression of ABCG2 (PX plus carbachol P=0.0270; PX plus APE P=0.0425 vs. control). The addition of carbachol, APE or PX alone did not modify the expression of this transporter in tumor cells (Figs 6A and 6B).

**Fig 6.**
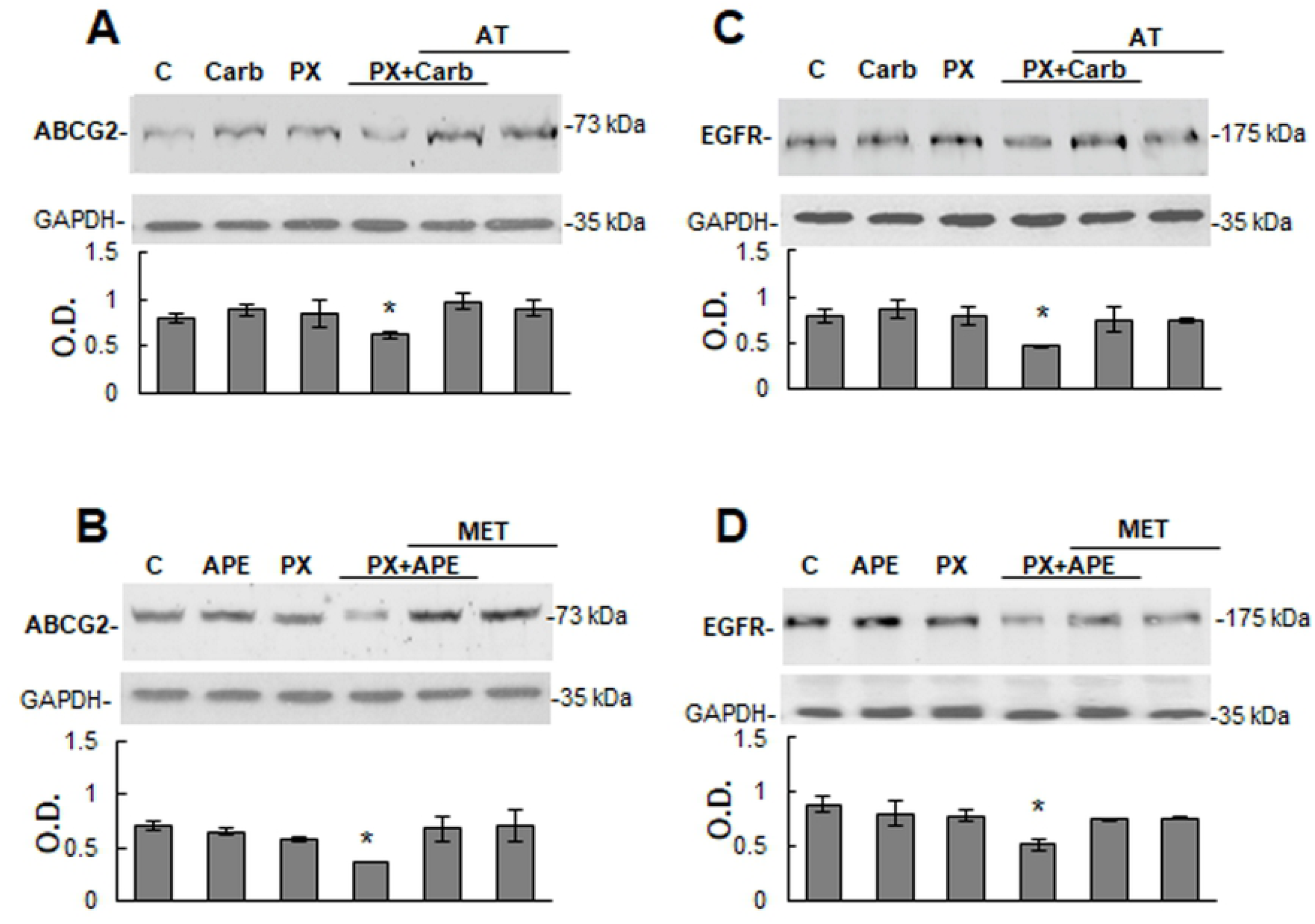
Expression of ATP binding cassette G2 transporter and epidermal growth factor receptor in MDA-MB231 cells. The expression of ATP binding cassette G2 (ABCG2) and epidermal growth factor receptor (EGFR) in tumor cells was analyzed by Western blot. Cells were treated for three cycles with paclitaxel (PX) (10^−8^M) combined with A) and C) carbachol (Carb) (8.6×10^−12^M) or with B) and D) arecaidine propargyl ester (APE) (1.1×10^−5^M) in the absence or presence of atropine (AT) (10^−9^M) or methoctramine (MET) (10^−5^M). Molecular weights are indicated on the right. Densitometric analysis of the bands was expressed as optical density (O.D.) units relative to the expression of glyceraldehyde 3-phosphate dehydrogenase (GAPDH) protein used as loading control. One representative experiment of 3 is shown (*P<0.05 vs. control).

It has been documented that the constitutively activation of EGFR or its transactivation contributes to drug resistance in different types of tumor cells (26). As it is shown in figure 6C, MDA-MB231 cells express this receptor and the addition of three cycles of PX plus carbachol significantly reduced protein expression by more than 40% in comparison to control (P=0.0477) (Fig 6C). Similarly, Western blot analysis showed that the treatment with APE combined with PX also down-regulated EGFR protein by more than 41% respect to control (P=0.0460) (Fig 6D).

### Effect produced by the combined treatment of muscarinic agonists and paclitaxel on MDA-MB231 cell migration

Since invasion is an important step in tumor progression, we analyzed the effect of PX plus carbachol or APE on tumor cell migration in an *in vitro* wound healing assay (Fig 7). At the end of experimental time (28 h) control wound was covered by 79±8% while the addition of PX plus carbachol or APE prevented wound covering by 38±5% or 47±6% respectively. Both effects were reverted in the presence of atropine or methoctramine.

**Fig 7.**
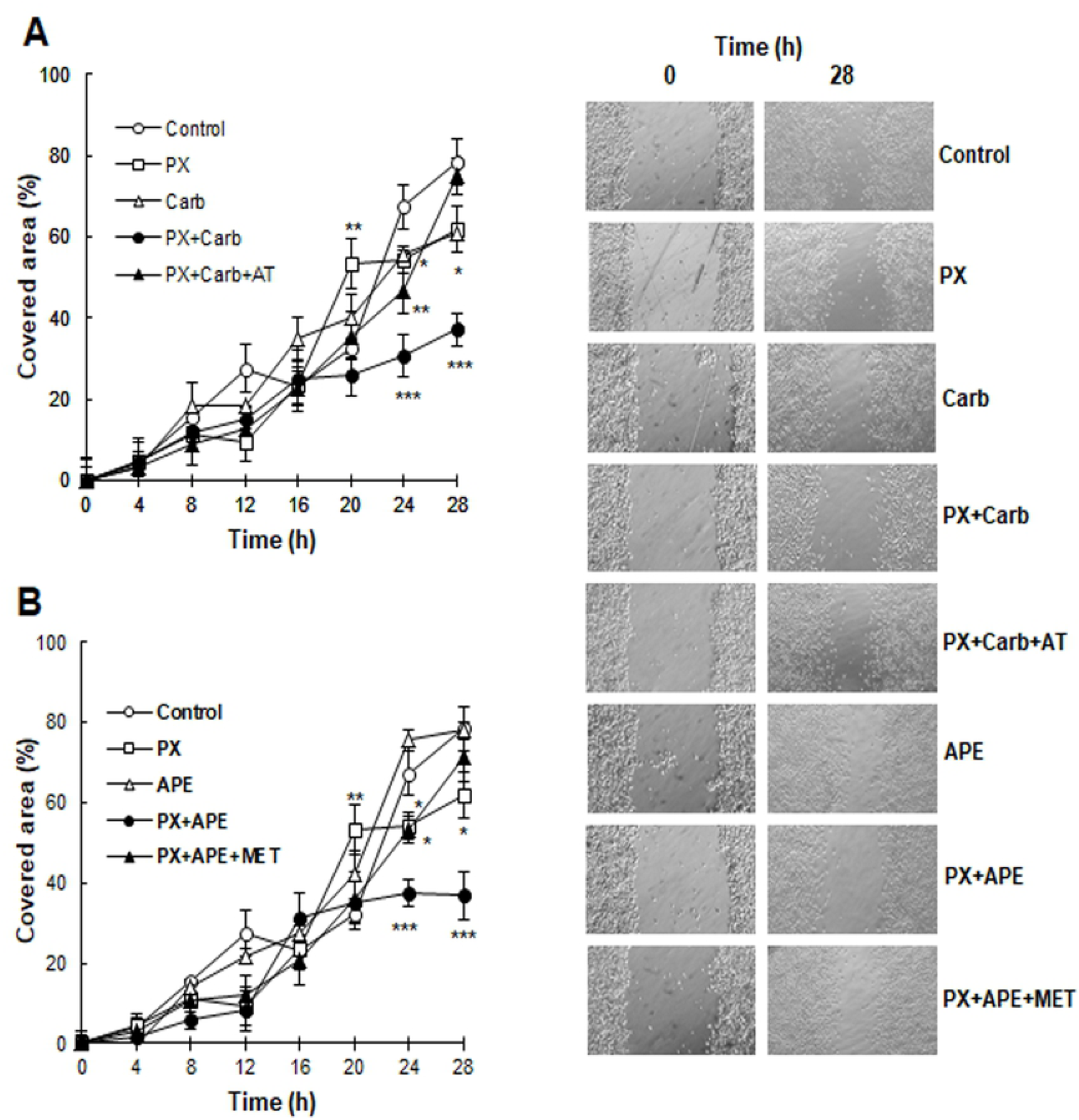
Effect of the combination of paclitaxel with a muscarinic agonist on MDA-MB231 cell motility. Cells were treated with A) paclitaxel (PX) (10^−8^M) combined with carbachol (Carb) (8.6×10^−12^M) in the absence or presence of atropine (AT) (10^−9^M) or PX (10^−8^M) was combined with arecaidine propargyl ester (APE) (1.1×10^−5^M) in the absence or presence of methoctramine (MET) (10^−5^M). Representative photographs (64X) were obtained by phase contrast at the beginning (0h) and at the end of experimental time (28 h). Values are mean ± S.E.M. of 3 experiments performed in duplicate. (*P<0.05; **P<0.01; ***P<0.0001 vs. control).

### The combined treatment reduces vascular endothelial growth factor-A and tumor induced-angiogenesis

Taking into account that metronomic administration of drugs can exert additional benefits in comparison to traditional chemotherapy, we analyzed the ability of both drug combinations to exerted anti-angiogenic actions. The addition of PX plus carbachol to MDA-MB231 cells in culture for three cycles significantly down-regulated VEGF-A expression by almost 39% (P=0.0490) (Fig 8A). Moreover, the addition of APE to the combination exerted a similar reduction in VEGF-A expression in MDA-MB231 cells (37±3%) (P=0.0257) (Fig 8B). We also analyzed the ability of MDA-MB231 cells to induce blood vessel formation in the skin of NUDE mice (Fig 8C). The inoculation of tumor cells increased by 31±5% (P=0.0362) skin neovascularization in comparison to sham skin. The i.p. administration of carbachol plus PX at metronomic doses to tumor bearers for three cycles potently reduced tumor induced-angiogenesis. This effect was reverted when animals were treated i.p. with the non-selective antagonist atropine previously to the combination. Also, APE combined with PX at low doses significantly reduced tumor neovascularization and the pre-treatment of tumor bearers with methoctramine, a selective M_2_ antagonist, prevented the latter effect. Neovascular response in the skin of tumor bearing mice treated with PX plus carbachol or APE did not differ from the vascular density exhibited by skin obtained from sham animals (3.07±0.10). Representative photographs of sham skin, positive neovascularized skin induced by MDA-MB231 cells (Control) and the inhibition of angiogenic response produced by the *in vivo* treatment of NUDE mice with PX plus carbachol or APE are shown in figure 8D.

**Fig 8.**
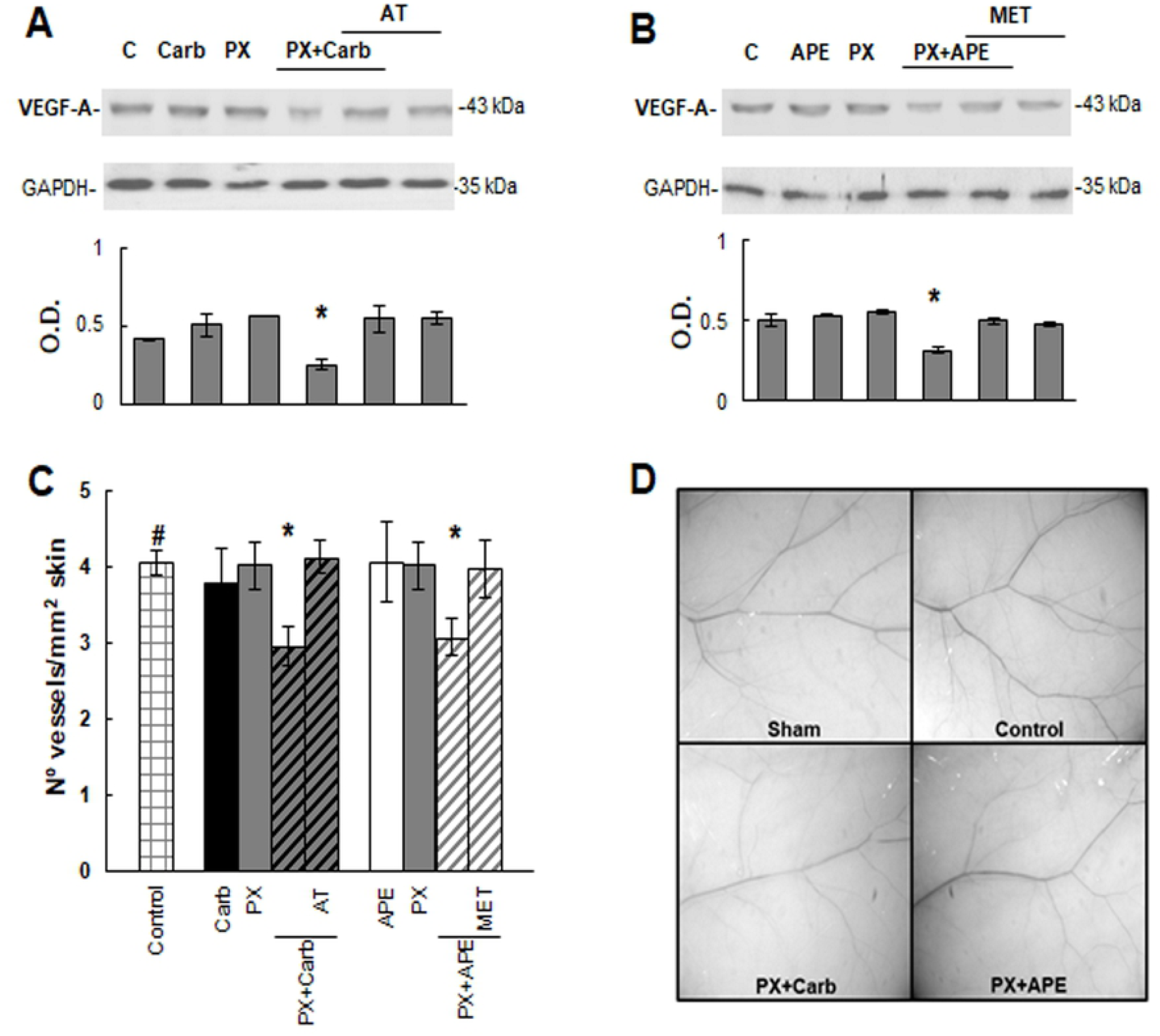
Tumor induced angiogenesis. To analyze the expression of vascular endothelial growth factor-A (VEGF-A) by Western blot, MDA-MB231 cells were treated with A) paclitaxel (PX) (10^−8^M) combined with carbachol (Carb) (8.6×10^−12^M) or B) with arecaidine propargyl ester (APE) (1.1×10^−5^M) in the absence or presence of atropine (AT) (10^−9^M) or methoctramine (MET) (10^−5^M) respectively. Molecular weights are indicated on the right. Densitometric analysis of the bands was expressed as optical density (O.D.) units relative to the expression of glyceraldehyde 3-phosphate dehydrogenase (GAPDH) protein used as loading control. One representative experiment of 3 is shown. C) *In vivo* neovascularization induced by MDA-MB231 cells in NUDE mice skin. Cells were inoculated as it was stated in Methods, and drugs were administered i.p. Values are mean ± S.E.M. of 3 experiments performed with 3 animals per group inoculated in both flanks. D) Representative photographs of mice skin from sham animals or inoculated with tumor cells (Control) without treatment or treated with PX+Carb or PX+APE. Magnification 6.4X. (#P<0.05 vs. sham skin; *P<0.05 vs. control).

## Discussion and conclusions

In this work, we demonstrated for the first time, the expression of different muscarinic receptor subtypes (1, 2, 4 and 5) in MDA-MB231 cells derived from a human TN mammary adenocarcinoma. We also observed that the long term addition either of carbachol or APE (non-selective and M_2_ receptor selective agonists, respectively) reduced cell viability in these breast tumor cells. Previous results from our group and also from other authors pointed to the ability of these muscarinic agonists to produce cell death in murine breast, bladder and neuronal tumor cells [14-16]. These results let us consider muscarinic receptors as specific therapeutic targets for the treatment of TN breast tumors since they are classified as very aggressive and malignant. Previous reports indicated that TN tumor bearers benefit from the addition of PX in the adjuvant therapy, supporting the conclusion that taxanes are useful for this group of patients [27]. Our results confirmed the ability of PX to decrease TN breast tumor cell viability, but it affects simultaneously non-tumorigenic MCF-10A breast cells as a side effect. The latter is one of the reasons because of specific adjuvant regimens for TN breast cancer are still under revision, and third generation chemotherapy regimens utilizing dose dense or metronomic poly-chemotherapy are thought to be more effective and less harmful than traditional chemotherapy. Considering the latter, together with our previous findings in MCF-7 cells that were sensible to a low dose metronomic therapy focused on muscarinic receptors [13]. Here, we designed a metronomic administration of PX plus muscarinic agonists at low concentrations. This strategy, besides focusing on muscarinic receptors as novel targets for the treatment of TN tumors, could be less aggressive for normal mammary cells that lack of these receptors [10]. Moreover, our results reveal the possibility of generalizing the usage of this combination of drugs since it is also effective when PX is replaced by doxorubicin. It is also important to consider that PX plus carbachol or APE is effective on MDA-MB468 cells, derived from a different TN tumor classified as Basal A with amplified EGFR [28].

It is important to note that M_2_ receptor mediates the main cytotoxic action of this low dose combined therapy, since similar results were obtained by replacing carbachol with APE, a selective M_2_ agonist, in the combination. The participation of this receptor subtype was confirmed by reversing the effect either with the preferential selective M_2_ antagonist methoctramine. In line with our findings, similar effects were demonstrated in human glioblastoma cell lines and in glioblastoma cancer stem cells [16,17]. As it can be seen from the results obtained in the current work, the presence of M_2_ receptors could be useful to target them with metronomic therapy in TN breast cancer treatment. However, this condition is important but not essential to focus metronomic therapy, since we had previously documented that a similar combination of PX plus carbachol was effective to reduce MCF-7 cell viability [13]. These breast cancer cells derived from a human luminal-A tumor, lack M_2_ receptors and express M_3_ and M_4_ subtypes. The previous results and those of this work, increase the importance of muscarinic receptors to be considered *in totum* as a target for oncological therapy.

One of the most important side effects of traditional chemotherapy is the resistance to drugs. In fact, the most commonly observed mechanism conferring drug resistance in cancer cells is the over-expression of ABC transporters that mediates the efflux of endogenous and exogenous substances using energy provided by ATP hydrolysis. ABCG subfamily plays pivotal roles in the transport of anticancer drugs out of cells, generating the development of drug resistance [29]. Here we analyzed the expression of ABCG2 since this protein is expressed in CD44^+^/CD24^-^ stem cell population that is abundant in MDA-MB231 tumor. The ABCG2 expression was potently down-regulated by the combination of PX either with carbachol or with APE. In relation with our results, Farhana et al. reported that malignancy of colon cancer cells induced by bile acids promotes cancer stemness in colonic epithelial cells by up-regulating M_3_ receptor together with ABCG2 expression [30].

Several aggressive tumors present an overexpression of the HER2 marker considering them a possible target in cancer therapy [31,32]. A variant with high homology of this marker is the EGFR which is also over-expressed in cancer [33,34], and whose activity is related to the increment in proliferation and invasion, and can be modulated by muscarinic receptors [35]. Our results demonstrated that the combination of PX plus carbachol or APE drastically reduced EGFR expression in tumor cells and this effect should be beneficial in the treatment of this type of tumor. Recently, it has been described that the inhibition of EGFR activity in glioblastomas produces a decrement in drug resistance activity mediated by the ATP-dependent membrane transporter, ABC [36]. Considering the link previously stated between ABC transporters’ expression and EGFR in glioblastomas, EGFR should be considered when designing a treatment for TN tumors in breast cancer patients.

It should be important to consider that, the reduction in cell viability produced by the combination of PX with carbachol or APE could be related on one hand to a decrement in the number of cancer stem cells which are present at very high levels in TN tumors and are generally positive to ABCG2 transporter as it was previously reported [37,38]. On the other hand, either the over-expression/activation of EGFR leads to VEGF-A production in MDA-MB231 cells and is closely related to the proliferative behavior of breast cancer cells as well as tumor endothelial cells [39]. In light of this background, it is expected that the combination of drugs designed in this work reduces proliferation and angiogenesis as observed in our results obtained *in vitro* and *in vivo* respectively.

Invasion is one of the most important steps in tumor progression since it is linked to metastasis and aggressiveness that are usually observed in TN breast cancer patients. Our results show for the first time the ability of low dose combined therapy to reduce tumor cell migration targeting M_2_ receptor. In addition, angiogenesis has been broadly considered as a switch-on mechanism in tumor growth and metastasis. One of the additional benefits of metronomic therapy to be tested is its ability to reduce tumor-induced angiogenesis [39]. Here we confirmed that the treatment of TN tumor cells with the combination of PX plus muscarinic agonists is able to reduce the expression of VEGF-A, and also the *in vivo* neovascular response induced by tumor cells.

In conclusion, our results demonstrate that low doses therapy combining PX with a non-selective or selective M_2_ agonist could be a useful strategy to treat TN breast tumors. In particular, the treatment focused on M_2_ receptor appears as a new promising therapeutic target to counteract not only breast cancer cell survival, but also to reduce invasion and pathological neo-angiogenesis, suggesting a possible prevention in drug chemoresistance by reducing ABCG2 and EGFR expression.

## Acknowledgments

The authors want to thank Mr. Francisco Sanchez, Mr. Daniel Gonzalez and Vet. Marcela Vázquez for their excellent technical assistance, and to Mrs. Patricia Fernández for her excellent management of financial support. We also want to thank Dr. Fernanda Troncoso and Lic. Vanina Vacheta from INQUIFIB-CONICET for providing MDA-MB468 cell line. This work was granted by CONICET PIP 2015-2017, 201501 00239; ANPCyT PICT 2015-2017, 2396 and UBA UBACYT 2014-2017, 20020130100168BA.

## Supporting information

**S1 Fig. Muscarinic receptors’ expression in MDA-MB468 cells.** Western blot assay to detect muscarinic (M) receptor subtypes. Molecular weights are indicated on the right. The expression of glyceraldehyde 3-phosphate dehydrogenase (GAPDH) protein was used as loading control. One representative experiment of 3 is shown.

